# Comparison of the 2^-C_T_^ method and the 2^-ΔΔC_T_^ method for real-time qPCR data analysis

**DOI:** 10.1101/2025.07.16.665089

**Authors:** Feng Lixiang, Zhao Rongqian, Kui Zhang, Wenxing Yang

## Abstract

**[purpose]:** The 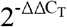 method proposed by Livak and Schmittgen in 2001 is used for real-time quantitative polymerase chain reaction (RT-qPCR) data analysis. This method’s fundamental logic involves normalizing all data against the control group to achieve relative quantification of gene expression. During our practical calculations, we identified an inherent bias in this method: the practice of directly taking arithmetic means of a set of C_T_ (or ΔC) values during 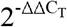 calculations fails to fully account for the exponential nature of C_T_ values (where C_T_ values serve as exponents in power-of-2 operations), leading to deviations in computational results. In this study, we elucidate this systematic bias and propose an innovative approach to circumvent it.

**[Methods]:** We propose the 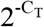 method, which bases calculations on 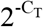 values and computes the fold-change of target genes relative to control group target genes. The computational logic of the 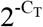 method strictly adheres to the exponential characteristics of C_T_ values, avoiding the use of arithmetic averaging at C_T_ and ΔC_T_ levels, thereby more accurately reflecting gene expression levels in datasets. In this paper, we detail the computational process of the 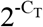 method and compare results from both methodologies.

**[Results]:** Calculations based on Livak and Schmittgen’s published data show differences between the two methods, though the discrepancies are relatively small. In calculations from our recent cadmium exposure experiments, the 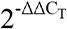 method indicates that 8-hour cadmium exposure increases *irg-6* gene expression in *C. elegans* from 1.314 to 7.125-fold, while the 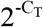 method shows an increase from 1 to 4.124-fold. The two methods exhibit nearly 70% discrepancy in *irg-6* gene fold-change quantification, with statistical p-values of 0.0002 and 0.0015 respectively. In all calculations presented in the paper, the 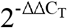 method fails to produce an exact unit value (mean fold-change ≠ 1) for control groups, whereas the 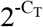 method consistently yields control group results of exactly 1, demonstrating its capability for precise normalization.

**[Conclusions]:** In summary, the 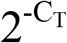 method demonstrates superior computational rigor, and we recommend its adoption in RT-qPCR data analysis. This improvement proves particularly valuable for experimental datasets with substantial C_T_ value variability among different samples, establishing a more reliable computational paradigm for RT-qPCR analysis.

## Introduction

In 2001, Livak and Schmittgen first proposed the comparative threshold cycle method (comparative C_T_ method, also known as the 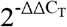 method) for analyzing real-time quantitative polymerase chain reaction (RT-qPCR) data to achieve relative quantification of gene expression [1]. In this method, the C_T_ value represents the number of cycles required for the RT-qPCR reaction to reach the preset fluorescence threshold, 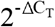 represents the normalized expression level of the target gene relative to the reference gene, and the 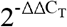 value represents the relative expression fold change of the target gene in the test sample. The basic logic of the 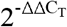 method is to compare the expression changes of the target gene based on the 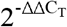 values of the target gene in each sample. According to Google Scholar statistics, as of May 21, 2025, this paper [1] has been cited nearly 190,000 times by scientists. In 2008, the two authors published a detailed description of the calculation method of the 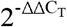 method in the journal *Nature Protocols* [2], which has been cited over 28,000 times. Undoubtedly, the 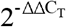 method is a widely recognized RT-qPCR data analysis method in the life sciences and medical fields and has been extensively used by the scientific community for over two decades.

The calculation of the 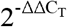 method essentially normalizes the expression level of the target gene in a specific sample relative to the control group [2]. Under this logic, we speculate that the mean 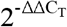 value of the control group should be the integer 1. However, neither in previously published papers using this method (e.g., Figure 4 in Assaei et al. [3], Figure 5 in Tan et al. [4], Figure 3 in Li et al. [5]) nor in the two highly cited papers by Schmittgen et al. [1, 2] is the mean 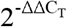 value of the control group the integer 1. These facts suggest that there may be an inherent bias in the calculation process of the 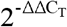 method. Through an in-depth analysis of the original calculation process of the 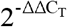 method, we found that in the actual calculation process [1, 2], the mean C_T_ value (C_T(mean)_) or mean ΔC_T_ value (ΔC_T(mean)_) is used to represent the C_T_ or ΔC_T_ values of a group of samples. This approach overlooks the exponential nature of C_T_ values (or ΔC_T_ values). This paper will report our findings from two aspects: first, we will explain the mathematical principle behind the bias in the calculation process of the 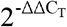 method; second, based on data from published papers and our cadmium exposure experiment data, we will fully describe the detailed calculation process of our proposed 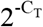 method.

## Materials and Methods

### Experimental Animals and Materials

The *Caenorhabditis elegans* used in this study were wild-type hermaphrodites of the N2 strain, purchased from the Caenorhabditis Genetics Center (USA). All nematodes were routinely cultured with *Escherichia coli* OP50 [6].

### Cadmium Exposure Experiment

When the nematodes reached adulthood, they were treated with alkaline sodium hypochlorite to collect eggs for synchronization. The collected eggs were cultured at a density of 200–300 nematodes per plate on nematode growth medium (NGM) plates [6] until the first day of adulthood, then divided into two groups (cadmium exposure group and control group) for the following experiments. Cadmium exposure group: nematodes were washed off the plates with 1 mL of M9 buffer and transferred to a 1.5 mL centrifuge tube. After natural sedimentation for 3 minutes, the supernatant was removed, and the nematodes were transferred to plates containing 100 μM cadmium chloride (CdCl□) for 8 hours of exposure. After exposure, the nematodes were washed three times with M9 buffer to remove residual bacteria, dehydrated with 70% ethanol for 5 minutes, and stored at −80°C for later use. Control group: after the initial washing and sedimentation, the nematodes were transferred to plates without CdCl□, with all other operations the same as the cadmium exposure group.

### RT-qPCR and Related Calculations

Total RNA was extracted from the collected nematode pellets using a column-based and Trizol method (R0027, Beyotime Biotechnology Co., Ltd.), and RNA concentration and purity were measured using a NanoDrop One spectrophotometer (Thermo Fisher Scientific Inc.). For each sample, 500 ng of total RNA was used to prepare cDNA libraries using the FastKing gDNA Dispelling RT SuperMix kit (KR220511, Tiangen Biotech Co., Ltd.). RT-qPCR was performed using the Universal SYBR qPCR Master Mix kit (BL697A, Bio-Rad Laboratories, Inc.) and a Bio-Rad CFX Connect real-time PCR system (Bio-Rad Laboratories, Inc.). The DNA sequences of the primers used in the experiment are as follows. Specific data analysis methods based on C_T_ values are described in the main text, and all calculation formulas have been embedded in the supplementary Excel files (Supplementary Tables 1 and 2). The supplementary tables are also available on Figshare.com [7].

Target gene irg-6:

5’-CTCAGTTCCAGACGTAGATACT-3’

5’-TCAATGTCTCCAGCAGCCTT-3’

Reference gene actin:

Forward: tctcttccagccatccttcttg

Reverse: tctgcatacgatcggcgatt

### Statistical Analysis

Data in this paper are expressed as mean ± SEM. Statistical analysis was performed using GraphPad Prism 10 software. A P-value < 0.05 was considered statistically significant.

## Results

### Calculation Logic of the 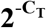 Method

C_T_ refers to the number of cycles required for the fluorescence signal in a PCR reaction to reach the set threshold. In the 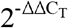 method, there are two scenarios where averaging at the C_T_ value level is required: ⍰ Using the arithmetic mean of C_T_ values from PCR replicates (C_T(mean)_) to represent the expression level of a specific gene in a sample; ⍰ Using the arithmetic mean of ΔC_T_ values (ΔC_T(mean)_) from all samples in a specific group to represent the relative expression level of a particular gene in that group. The calculation methods for C_T(mean)_ and ΔC_T(mean)_ are shown in Equations 1 and 2, respectively, where n represents the number of PCR replicates or samples in the group. This averaging logic is demonstrated in all five examples provided by Schmittgen et al. in Nature Protocols [1]. Taking PCR technical replicates as an example, when calculating the average expression level of gene A across n replicates, the 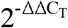 method directly takes the arithmetic mean of the C_T_ values from each replicate (Equation 1). This linear averaging approach ignores the exponential growth characteristic of PCR amplification.

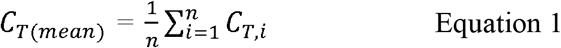

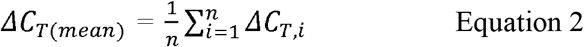

According to the fundamental principle of RT-qPCR, we generally assume that the amplification efficiency (E) is the same for all genes. Let the initial expression level of gene A in a sample be A_o_, and the quantity of gene A after C_T_ PCR cycles be K_T_. The relationship between C_T_, E, A_o_, and K_T_ can be expressed by Equation 3, which shows that CT values have an exponential nature. Equation 3 can be equivalently transformed into Equation 4. Since K_T_ is a constant, Equation 4 indicates that (1+E)^−CT^ is linearly and positively correlated with the initial expression level A_o_ of gene A (Equation 5).

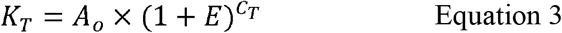

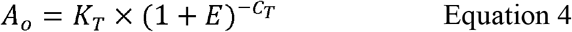

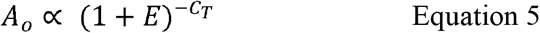

Based on the second assumption of the 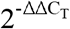 method principle [1], which states that the PCR amplification efficiency (E) equals 100% (in this case, 1 + E = 2), Equations 4 and 5 can be simplified to Equations 6 and 7, respectively. From Equation 7, it can be seen that 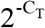 can serve as a linear representation of A_o_.

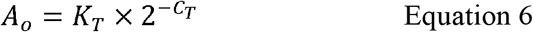

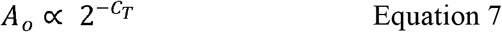

In the 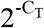 method, we first calculate the 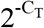 values for the target gene and the reference gene in each replicate, denoted as 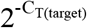 and 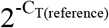, respectively. Then, the mean 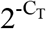 values of all replicates for a specific sample are used to represent its relative initial expression levels, denoted as T_o_ and R_o_ (Equations 8 and 9, where “n” represents the number of PCR replicates). Thus, the relative expression level (E_T_) of the target gene to the reference gene in a given sample can be calculated as “T_o_/R_o_” (Equation 10).

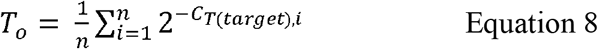

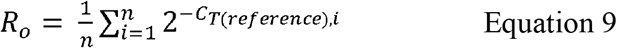

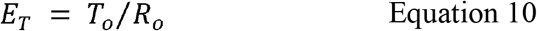

The mean E_T_ of the target gene in the control group E_T(control)_ is calculated based on all control samples (Equation 11, where “n” represents the number of samples in the control group). All data are normalized based on E_T(control)_ to obtain the fold change expression level A_FC_ of the target gene in all samples (Equation 12).

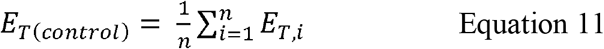

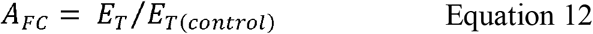

Finally, statistical analysis is performed based on A_FC_ to compare the differences in target gene expression levels between different groups. We named this method for comparing gene expression differences the 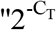 method.” Table 1 summarizes the formulas involved in the calculation processes of both the 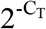 method and the 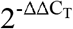 method. As can be seen from the equations, the 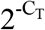 method respects the exponential nature of C_T_ values by performing calculations at the exponential level after transforming the C_T_ values. In terms of relative quantification of gene expression levels, the 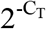 method is more accurate than the 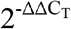 method.

**Table 1.**
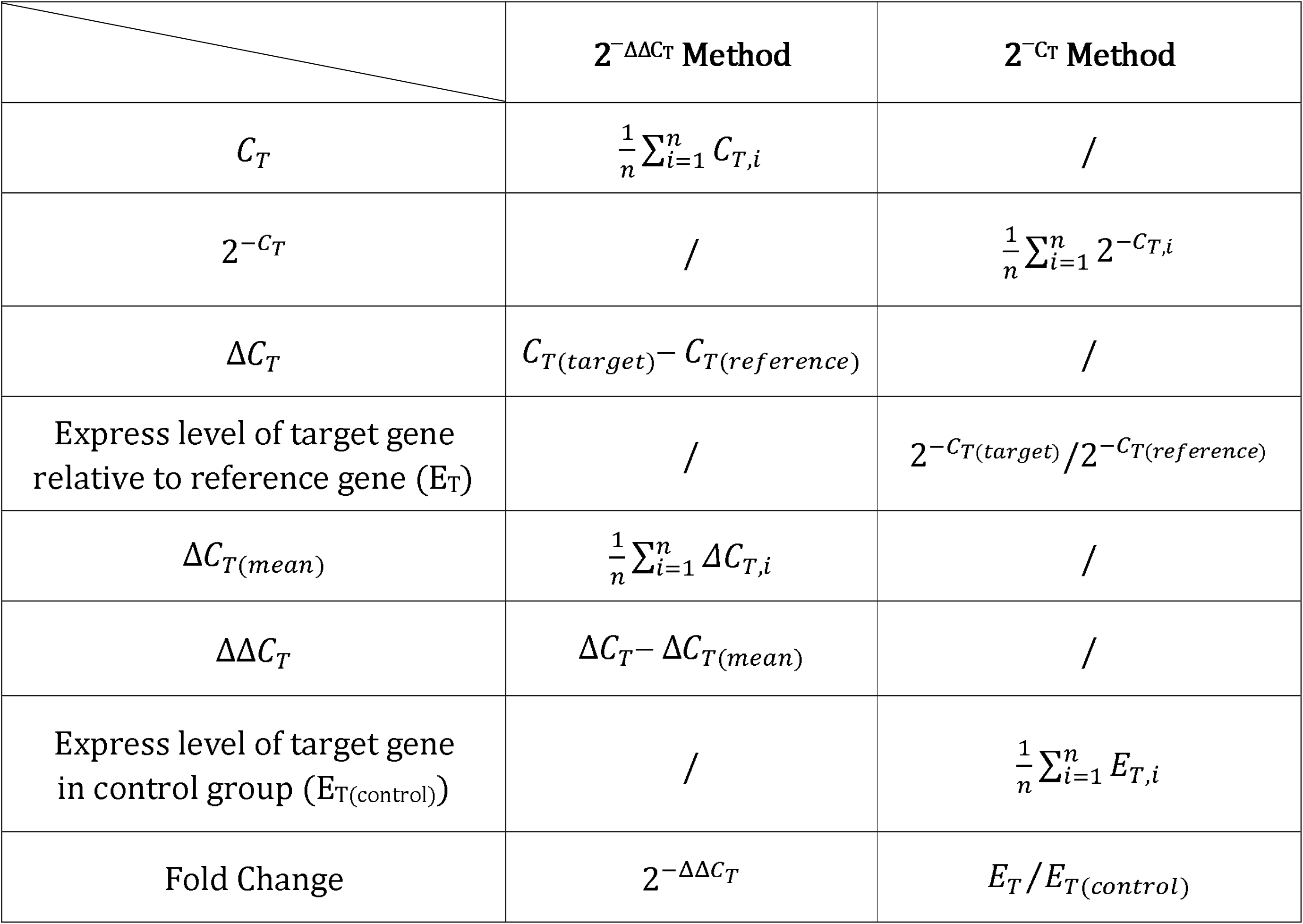
Comparisons of Calculation formula of both method

|  | **$2^{-\Delta\Delta C_T}$ Method** | **$2^{-C_T}$ Method** |
| --- | --- | --- |
| $C_T$ | $\frac{1}{n} \sum_{i=1}^n C_{T,i}$ | / |
| $2^{-C_T}$ | / | $\frac{1}{n} \sum_{i=1}^n 2^{-C_{T,i}}$ |
| $\Delta C_T$ | $C_{T(target)} - C_{T(reference)}$ | / |
| Express level of target gene<br>relative to reference gene ($E_T$) | / | $2^{-C_{T(target)}} / 2^{-C_{T(reference)}}$ |
| $\Delta C_{T(mean)}$ | $\frac{1}{n} \sum_{i=1}^n \Delta C_{T,i}$ | / |
| $\Delta\Delta C_T$ | $\Delta C_T - \Delta C_{T(mean)}$ | / |
| Express level of target gene<br>in control group ($E_{T(control)}$) | / | $\frac{1}{n} \sum_{i=1}^n E_{T,i}$ |
| Fold Change | $2^{-\Delta\Delta C_T}$ | $E_T / E_{T(control)}$ |

### Comparison of results from analyzing published data using both methods

In their original paper introducing the 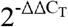 method [1], Schmittgen’s team presented their RT-qPCR experimental results for the target gene fos-glo-myc and reference gene actin. The data in the gray cells of Table 2 and Supplementary Table 1 were sourced from Livak and Schmittgen’s paper [1], while the remaining data were calculated by us using the 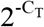 method. Based on Equation 1, the C_T(mean)_ for fos-glo-myc was 21.9, and for actin it was 22.5; based on Equation 2, the ΔC_T(mean)_ for fos-glo-myc relative to actin was -0.6 (Supplementary Table 1). After normalization, the expression levels of fos-glo-myc were 1.023, 0.786, and 1.243, with a mean expression level of 1.02 (Table 2). According to the 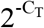 method calculations, the expression levels of fos-glo-myc were 0.986, 0.801, and 1.214, with a mean expression level of exactly 1.000, demonstrating that the 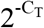 method can indeed perfectly normalize the control group data. To fully present the detailed calculation process for all data, we summarized the calculation procedures for all data in Table 2 in Supplementary Table 1.

**Table 2.**
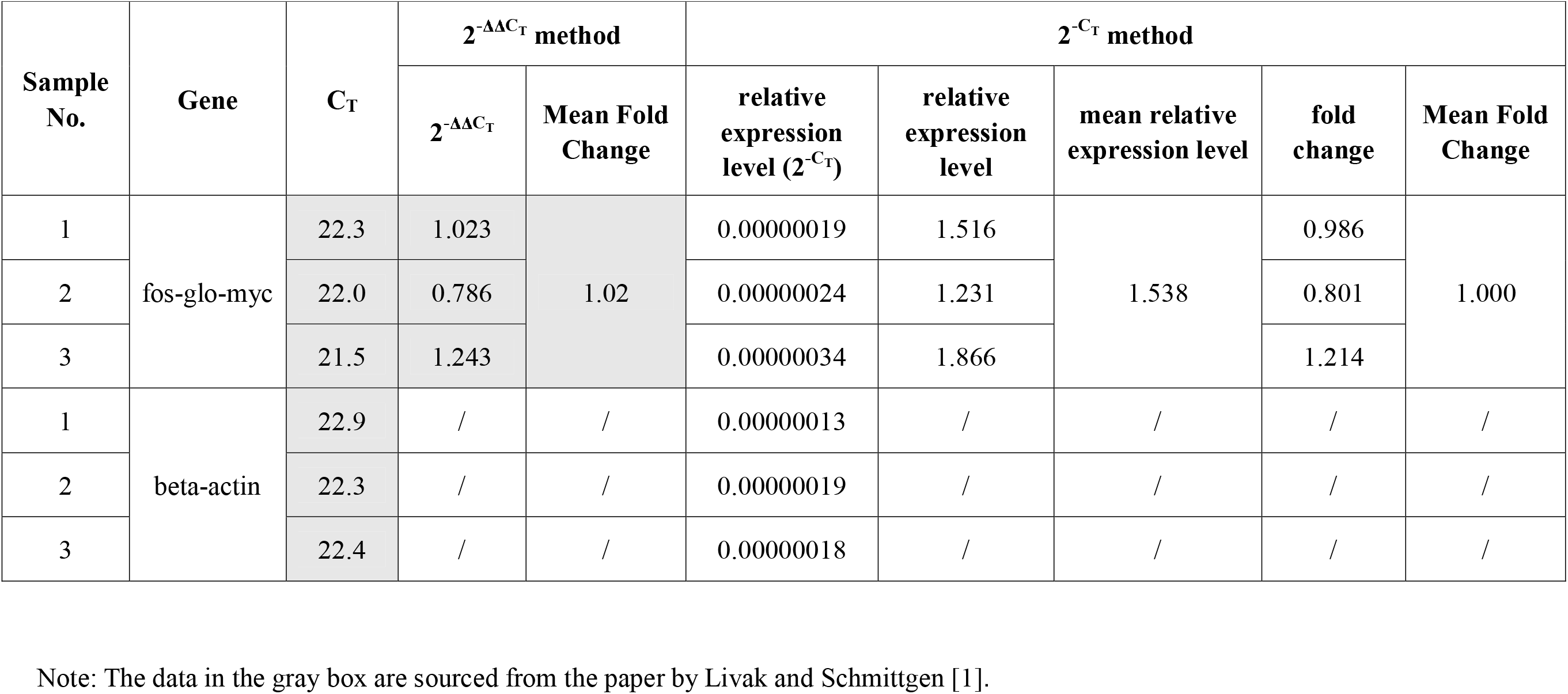
Calculations following the Schmittgen method

### Analysis of cadmium exposure effects on irg-6 gene expression in C. elegans using both 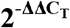 and 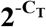 methods

Based on Livak and Schmittgen’s paper [1], we were unable to fully present the specific calculation process related to PCR replicate data. Therefore, we systematically compared the differences in calculation results between the 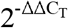 and 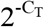 methods using data from our laboratory’s recent cadmium exposure experiment.

We exposed adult C. elegans to cadmium for 8 hours to observe its effects on irg-6 gene expression levels. For each sample, we performed RT-qPCR with three replicates. In the replicate-related calculations, the 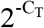 method calculated the arithmetic mean of the exponential-transformed C_T_ values for replicates based on Equations 8 and 9 (Supplementary Table 2). Then, using Equations 10-12, we calculated the irg-6 gene expression levels in the control group as 0.749, 2.259, 0.832, and 0.160, with a mean of exactly 1; the irg-6 gene expression levels in the cadmium-exposed group were 4.473, 5.167, 3.567, and 3.648, with a mean of 4.214 (Table 3 and Supplementary Table 2).

**Table 3.** The impact of cadmium exposure on *irg-6* gene expression levels in *Caenorhabditis elegans* calculated by 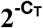 Method

| Group | Gene name | sample No. | mean relative expression level (mean of $2^{-C_T}$) | relative expression level of target gene to reference gene | mean relative expression level of target gene | fold change | mean fold change |
| --- | --- | --- | --- | --- | --- | --- | --- |
| control | irg-6 | 1 | 0.000000000032 | 0.0000301 | 0.0000402 | 0.749 | 1.000 |
|  |  | 2 | 0.000000000105 | 0.0000907 |  | 2.259 |  |
|  |  | 3 | 0.000000000352 | 0.0000334 |  | 0.832 |  |
|  |  | 4 | 0.000000000098 | 0.0000064 |  | 0.160 |  |
|  | actin | 1 | 0.000001068055 | / | / | / | / |
|  |  | 2 | 0.000001161385 | / | / | / | / |
|  |  | 3 | 0.000010535024 | / | / | / | / |
|  |  | 4 | 0.000015145803 | / | / | / | / |
| Cd-treated | irg-6 | 1 | 0.000000000227 | 0.0001796 | / | 4.473 | 4.214 |
|  |  | 2 | 0.000000000303 | 0.0002075 | / | 5.167 |  |
|  |  | 3 | 0.000000000999 | 0.0001432 | / | 3.567 |  |
|  |  | 4 | 0.000000001518 | 0.0001465 | / | 3.648 |  |
|  | actin | 1 | 0.000001262155 | / | / | / | / |
|  |  | 2 | 0.000001460077 | / | / | / | / |
|  |  | 3 | 0.000006978325 | / | / | / | / |
|  |  | 4 | 0.000010365774 | / | / | / | / |

Table 4 and Supplementary Table 2 present our analysis of irg-6 gene expression changes after 8-hour cadmium exposure using the 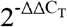 method. Based on Equation 1, the C_T(mean)_ for irg-16 in the control group was 33.467, and for actin it was 18.031; based on Equation 2, the ΔC_T(mean)_ for irg-6 relative to actin was 15.436 (Table 4). After normalization, the mean fold change of irg-6 expression was 1.314, again demonstrating that the 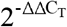 method struggles to normalize control group data to exactly 1. Additionally, the irg-6 expression level in the cadmium-exposed group was 7.125-fold higher than in the control group (Table 4 and Supplementary Table 2). Compared to the 4.214-fold change calculated by the 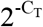 method, the 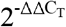 method’s result was nearly 70% higher. This difference indicates that the calculation results of the two methods may show significant discrepancies in specific datasets.

**Table 4.** The impact of cadmium exposure on *irg-6* gene expression levels in *Caenorhabditis elegans* calculated by 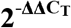 Method

| Group | Gene name | sample No. | $C_{T(\text{mean})}$ | $\Delta C_T$ | $\Delta C_{T(\text{mean})}$ | $\Delta\Delta C_T$ | $2^{-\Delta\Delta C_T}$ | Mean Fold Change |
| --- | --- | --- | --- | --- | --- | --- | --- | --- |
| control | irg-6 | 1 | 35.05 | 15.210 | 15.436 | -0.226 | 1.169 | 1.314 |
|  |  | 2 | 33.73 | 14.007 |  | -1.429 | 2.693 |  |
|  |  | 3 | 31.82 | 15.287 |  | -0.149 | 1.109 |  |
|  |  | 4 | 33.26 | 17.240 |  | 1.804 | 0.286 |  |
|  | actin | 1 | 19.84 | / | / | / | / | / |
|  |  | 2 | 19.72 | / | / | / | / | / |
|  |  | 3 | 16.54 | / | / | / | / | / |
|  |  | 4 | 16.02 | / | / | / | / | / |
| Cd-treated | irg-6 | 1 | 32.17 | 12.567 | / | -2.869 | 7.31 | 7.125 |
|  |  | 2 | 31.73 | 12.333 | / | -3.103 | 8.59 |  |
|  |  | 3 | 29.98 | 12.853 | / | -2.583 | 5.99 |  |
|  |  | 4 | 29.30 | 12.710 | / | -2.726 | 6.62 |  |
|  | actin | 1 | 19.60 | / | / | / | / | / |
|  |  | 2 | 19.40 | / | / | / | / | / |
|  |  | 3 | 17.13 | / | / | / | / | / |
|  |  | 4 | 16.59 | / | / | / | / | / |

We performed statistical analysis on the results from both methods and found that the fold changes calculated by both methods were statistically significant (Figure 1), indicating that 8-hour cadmium exposure significantly increased irg-6 gene expression. The p-value for the 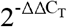 method’s results was 0.0002, while the p-value for the 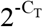 method’s results was 0.0015, showing an order of magnitude difference between them.

**Fig 1.**
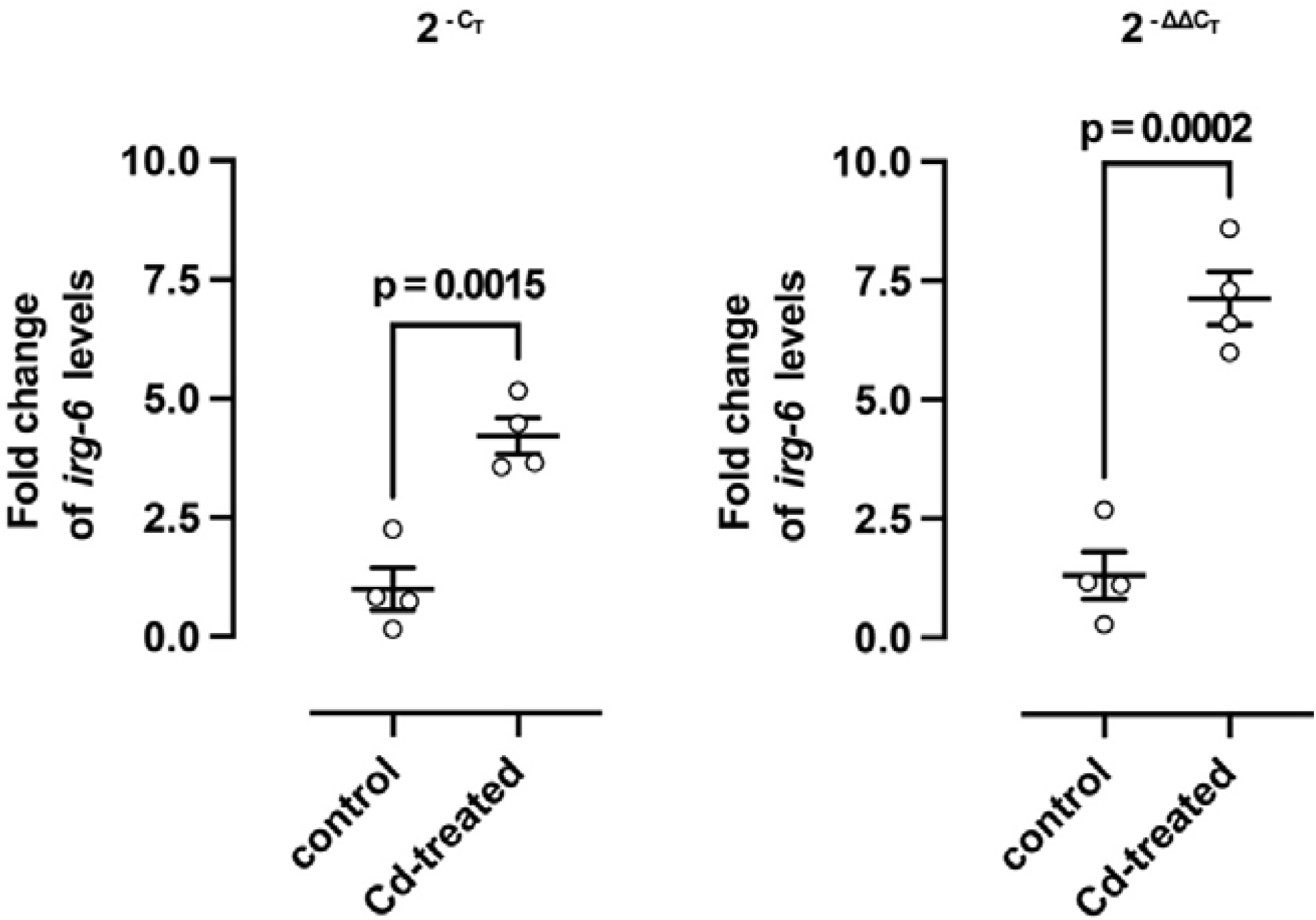
The effect of cadmium exposure on the expression of the *irg-6* gene in *Caenorhabditis elegans*. An independent samples *t* test was performed to compare the means between control and Cd-treated groups.

To evaluate the consistency between the 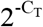 and 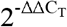 methods in expression level calculations, we performed Pearson correlation analysis on the fold change results obtained from both methods. The analysis showed an extremely high linear positive correlation between them (Figure 2, r=0.9909, R^2^=0.9819), but the values calculated by the 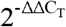 method were significantly higher than those from the 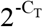 method. This suggests that although the two methods have different calculation logics and show high consistency in expression trend determination, there are systematic differences in the absolute values of fold changes, which is precisely the systematic bias issue we are concerned about.

**Fig 2.**
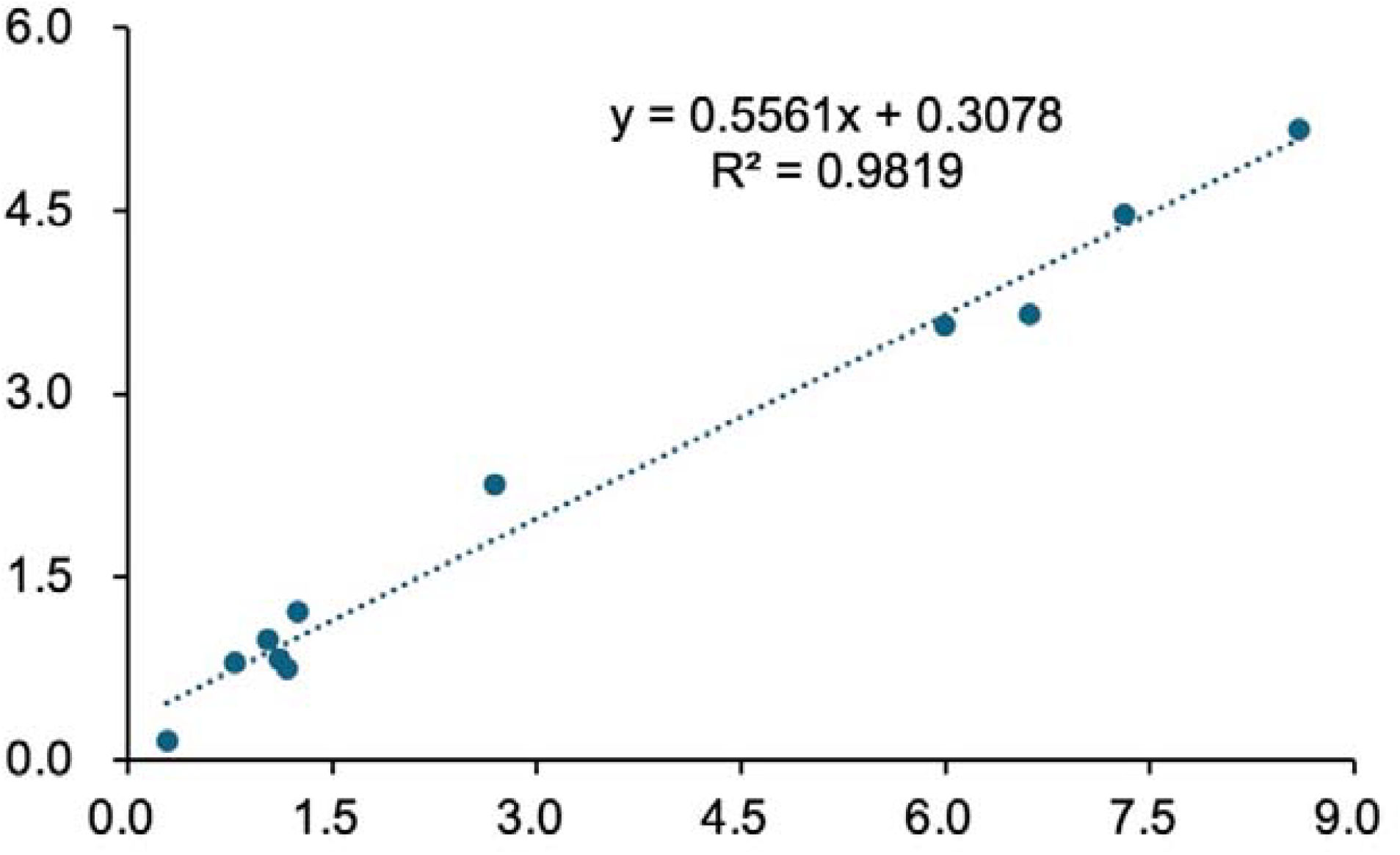
Pearson’s correlation analysis between the results calculated using the 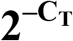 method and the 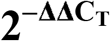 method. y axis, results calculated using the 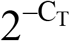 method; x axis, results calculated using the 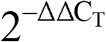 method.

### Impact of data dispersion on experimental results

We compared the dispersion levels between our cadmium exposure data and the data from Livak’s paper [1]. Since Livak’s paper had a small sample size (sample number=1, gene number=2), making variance significance testing inappropriate, we used descriptive statistics for comparison. As shown in Table 5, in our cadmium exposure data, the standard deviation (SD) of CT values for each sample ranged from 0.08 to 1.57, with an overall dispersion level (SD of SDs) of approximately 0.47. In Livak’s paper, the CT value SDs ranged from 0.32 to 0.4, with a dispersion level of about 0.06. These results suggest that our cadmium exposure data had greater overall fluctuations and higher inter-sample dispersion. The difference between the results calculated by the 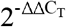 and 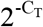 methods for this dataset was also relatively large (Tables 3 and 4, 7.125 vs. 4.214). This finding suggests that the greater the dispersion of the original data, the larger the potential discrepancy between the results of the two methods.

**Table 5.** Assessment of data variability

| group |  | gene | sample ID | mean C<sub>T</sub> | SD of C<sub>T</sub> | mean SD | SD of SD | Coefficient of Variation (%) |
| --- | --- | --- | --- | --- | --- | --- | --- | --- |
| our experimental data | control | irg-6 | 1 | 35.05 | 0.95 | 0.48 | 0.47 | 2.71 |
|  |  |  | 2 | 33.73 | 1.57 |  |  | 4.67 |
|  |  |  | 3 | 31.82 | 1.37 |  |  | 4.29 |
|  |  |  | 4 | 33.26 | 0.19 |  |  | 0.58 |
|  |  | actin | 1 | 19.84 | 0.12 |  |  | 0.61 |
|  |  |  | 2 | 19.72 | 0.18 |  |  | 0.93 |
|  |  |  | 3 | 16.54 | 0.10 |  |  | 0.60 |
|  |  |  | 4 | 16.02 | 0.23 |  |  | 1.44 |
|  | Cd-treated | irg-6 | 1 | 32.17 | 0.76 |  |  | 2.37 |
|  |  |  | 2 | 31.73 | 0.67 |  |  | 2.10 |
|  |  |  | 3 | 29.98 | 0.62 |  |  | 2.08 |
|  |  |  | 4 | 29.30 | 0.15 |  |  | 0.50 |
|  |  | actin | 1 | 19.60 | 0.14 |  |  | 0.70 |
|  |  |  | 2 | 19.40 | 0.22 |  |  | 1.13 |
|  |  |  | 3 | 17.13 | 0.08 |  |  | 0.44 |
|  |  |  | 4 | 16.59 | 0.38 |  |  | 2.28 |
| data from Livak paper |  | fos-glo-myc | 1 | 21.93 | 0.40 | 0.36 | 0.06 | 1.84 |
|  |  | beta-actin | 1 | 22.53 | 0.32 |  |  | 1.43 |

Note: SD, standard deviation; coefficient of variation = $\frac{SD \text{ of } C_T}{mean C_T} \times 100\%$ [8].

According to Taylor et al.’s systematic review [8], a coefficient of variation (CV) of less than 10% for technical replicates per sample indicates good qPCR reproducibility. As shown in Table 5, all data in our cadmium exposure experiment had CVs below 10%. This result indicates that although the cadmium exposure data showed high dispersion levels, the experimental reproducibility was good, and operational errors were minimal. Variability in qPCR CT values can arise from multiple factors, including biological differences, sample composition, RNA extraction efficiency, and sampling errors, which do not necessarily reflect unreliable experimental operations [8]. In our experiment, using pooled samples (whole populations of worms), the CT variability partially reflected the true biological differences in individual gene expression.

## Discussion

In this study, we systematically examined the inherent bias in the 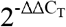 method proposed by Livak and Schmittgen in 2001 for RT-qPCR data analysis. In their original paper, the mathematical derivation of the 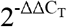 method was based on C_T_ values from a single sample containing both target and reference genes. Thus, the theoretical framework did not account for averaging operations. However, in practical applications - whether calculating representative C_T(mean)_ values from technical replicates or determining ΔC_T(mean)_values across multiple control samples - the 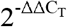 method employs arithmetic averaging, which disregards the exponential nature of C_T_ values.

Our formula analysis reveals fundamental differences between the calculation logics of the 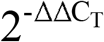 and 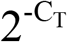 methods, leading to consistently divergent results. This discrepancy was confirmed through both our re-analysis of published data and new cadmium exposure experiments. While the deviation was relatively small (1.02 vs 1.00) in the published dataset, it reached 70% in our cadmium exposure study, with nearly an order-of-magnitude difference in statistical p-values. We hypothesize that this variation correlates with data dispersion - greater dispersion in raw data may amplify differences between the two methods. Consequently, statistically borderline results (e.g., p≈0.05) might lose significance when calculated using the alternative method. Based on mathematical rigor, we conclude that the 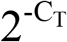 method represents a more theoretically sound approach and recommend its adoption for RT-qPCR data analysis.

Our derivation assumes 100% PCR amplification efficiency (E=1, yielding doubling amplification). While actual reactions rarely achieve perfect efficiency (typically E<1), the fundamental relationship 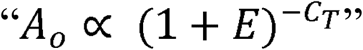 (Equation 5) remains valid. The exponential nature of C_T_ values persists regardless of E, maintaining the necessity for exponential transformation before averaging. For E<1, simply replace “2” with “(1+E)” in all formulas. The core principle of the 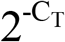 method - avoiding linear averaging of raw C_T_ values while operating on exponentially transformed data - retains universal applicability across different amplification efficiencies, ensuring broad methodological utility.

## Author Contribution

FENG Lixiang: Data collection and analysis, manuscript drafting. ZHAO Rongqian: Data validation. ZHANG Kui: experimental design. YANG Wenxing: experimental design, manuscript drafting, funding acquisition. All authors contribute to manuscript revision, discussion, and have approved the submission and agree to be accountable for all aspects of the work.

